# Pharmacologic manipulation of complement receptor 3 prevents dendritic spine loss and cognitive impairment after acute cranial radiation

**DOI:** 10.1101/2020.11.25.398701

**Authors:** Joshua J. Hinkle, John A. Olschowka, Jacqueline P. Williams, M. Kerry O’Banion

**Author notes:** **Correspondence:** M. Kerry O’Banion, 601 Elmwood Avenue, Box 603, Rochester, NY 14642.

## Abstract

Cranial irradiation induces healthy tissue damage that can lead to neurocognitive complications and negatively impact patient quality of life. One type of damage associated with cognitive impairment is loss of neuronal spine density. Based on developmental and disease studies implicating microglia and complement in dendritic spine loss, we hypothesized that irradiation-mediated spine loss is microglial complement receptor 3 (CR3)-dependent, and associated with late-delayed cognitive deficits. Utilizing a model of cranial irradiation (acute, 10 Gy gamma) in C57BL/6 mice we found that male mice demonstrate irradiation-mediated spine loss and cognitive deficits whereas female mice and CR3 knockout mice do not. Moreover, pharmacological blockade of CR3 with leukadherin-1 (LA1) prevented these changes in irradiated male mice. Interestingly, CR3 KO mice showed reduced behavioral task performance suggesting that CR3 is important for normal learning and memory. Improving our understanding of irradiation-mediated mechanisms and sexual dimorphic responses is essential for the identification of novel therapeutics to reduce irradiation-induced cognitive decline and improve patient quality of life.

## Introduction

Cranial radiation therapy is the leading mode of treatment for primary and metastatic malignancies in the brain with protocols that depend on several factors; including patient age, current comorbidities, tumor prognosis, and tumor burden (1,2). Unfortunately, cranial radiation is associated with significant late-delayed effects and depending on dose and irradiated tissue volume, as well as the brain region(s) affected, progressive, irreversible symptoms may manifest as physical weakness, changes in mood, and cognitive dysfunction (3,4). Eighty percent of patients that receive whole brain radiotherapy and survive more than six months develop some form of cognitive dysfunction which impacts their lives; 5% of these patients progress from impairment to dementia requiring 24 hour care (1). Other than reducing the radiation field to the point where it might compromise tumor treatment, there are no treatments to ameliorate injury to normal brain tissue and irradiation-mediated cognitive decline. Part of the difficulty is we do not fully understand the effects of radiation on normal tissue. While late-delayed brain symptoms have been associated with overt loss of white matter volume via necrosis, cognitive deficits can occur without gross alterations in brain anatomy (5), suggesting a subtler disruption of neural function.

Pre-clinical studies utilizing rodents have demonstrated hippocampal-dependent impairments in multiple behavioral tasks (6) suggesting that the hippocampus is an important target in irradiation-mediated cognitive decline, a result supported by hippocampal sparing radiotherapy strategies (7). The interconnectivity of the hippocampal network depends on the dynamic modification and remodeling of dendritic spines to allow synaptic plasticity and proper allocation of experience-dependent memories (8). Thus, alterations in dendritic spine formation and elimination may reflect changes in synaptic efficacy that are functionally related to learning and memory. These alterations may also underlie cognitive impairment in aging and neurologic disorders, which are associated with dendritic abnormalities and atypical spine densities (9). Following moderate doses of low-LET radiation, reductions in dendritic arbor complexity and spine density, as well as altered spine morphology have been reported (10,11) that likely compromise neural connectivity and memory function. Similar alterations have been observed following modest doses of protons and heavy ions (12,13). While several mechanisms may contribute to neuronal dysfunction, studies utilizing colony stimulating factor-1 receptor (CSF1R) inhibitors ameliorated irradiation-mediated deficits in hippocampal-dependent behavioral tasks (14-16) implicating microglia as possible effectors of radiation-induced injury and subsequent cognitive impairment.

Microglia, the resident immune macrophages of the central nervous system (CNS), rapidly respond to adverse stimuli in order to restore homeostatic function. Microglia also modulate neural networks through secretion of neurotrophic transmitters or directly through phagocytic maintenance (17). Indeed, microglia play a critical role in regulating synaptic and structural remodeling during development and plasticity (18). However, when an inflammatory response persists to a chronic state, microglial-dependent functions can shift from beneficial to destructive and exacerbate neuronal injury (19). Post-irradiation, microglia become reactive, secrete pro-inflammatory mediators, and upregulate phagocytic machinery (20). While an enhanced phagocytic state doesn’t directly implicate microglia in irradiation-mediated dendritic spine loss and cognitive decline, the aforementioned microglial depletion studies raise this possibility. However, a key piece is missing as cellular components require identification and recognition for microglial removal. Based on developmental, aging, injury, and disease models (21-24), we hypothesized that complement pathway activation could play a role in the irradiation injury response by linking increased inflammation and microglial reactivity to neuronal injury and dendritic spine loss.

The complement system comprises three pathways (classical, alternative, and lectin) that converge at C3. Once activated, C3 undergoes proteolytic cleavage into activating fragments C3a and opsonins C3b, iC3b, C3c. C3a is involved in inflammatory responses and microglial chemotaxis while C3b and iC3b are recognized by complement receptor-1 (CR1) and -3 (CR3), respectively, and lead to the phagocytosis of opsonin-tagged material (25). Microglia, as macrophages, express high levels of C1q and CR1, appear to exclusively express CR3 (CD11b/CD18 or mac-1) in the healthy brain (26), and upon activation, can upregulate C1q, CR1, C3, and CR3 in order to recognize, engulf, and remove opsonized material (27).

The C1q-dependent classical complement pathway has been implicated in CNS development with C1q and C3 localizing to weaker synapses and prompting microglial-mediated CR3-dependent phagocytosis (24,28). During aging, C1q and C3 are upregulated in mouse and human brains (29) and are associated with an increased pro-inflammatory cytokine profile and a decrease in synaptic transmission genes (30). Further, C3 deficiency enhanced hippocampal-dependent spatial learning in young and aging mice, suggesting that C3-dependent synapse loss is detrimental in aging (23,31). Similarly, complement is implicated in microglial-mediated synapse loss in injury response and disease progression. For instance, C1q, C3, or CR3 deficiency prevents spine loss and mitigates cognitive impairment in a mouse model of AD (32). Together, these studies provide insight into microglial-mediated spine loss and suggest a common mechanism that is complement-dependent and associated with cognitive dysfunction.

Interestingly, radiation injury shares gene profiles and molecular pathways with aging (33) and dementia (34) suggesting that complement may be involved; however, relatively few studies have explored the role of complement in irradiation-mediated brain injury. One study revealed that C3 KO mice had increased microglial numbers, higher acute levels of pro-inflammatory mediators (IL-1β and IL-6), and enhanced place learning capacity 3 months post-irradiation at 10 days of age (8 Gy) (35). Work by our group demonstrated microglial activation associated with a loss of spine density in male mice cranially-irradiated with 10 Gy that was mitigated in CR3 KO mice (36); however, whether mitigating irradiation-mediated spine loss is beneficial to cognitive function in the absence of CR3 remains unknown and is the premise for the current study.

To investigate cranial irradiation-mediated effects on neuron-glia interactions and cognitive function we focused on dendritic spine density and microglial reactivity in the hippocampus of male and female mice. To replicate previous findings (11) and improve upon our prior results (36), we crossed Thy1-YFP mice with CR3 WT and KO mice to ask whether CR3 deficiency can mitigate irradiation-mediated spine loss and cognitive dysfunction in behavioral tasks. Further, since CR3 is globally knocked-out and could affect developmental events, we used a commercially available CD11b agonist, Leukadherin-1 (LA1), to temporally modify CR3 signaling in Thy1-YFP::CR3 WT mice.

## Methods

### Animals

All animal procedures were carried out with ethical standards recommended by the Panel on Euthanasia of the American Veterinary Medical Association and approved by the University of Rochester Institutional Animal Care and Use Committee. Animal strains and conditions were as previously described (36) with the exception that *CR3*^+/+^ and *CR3*^-/-^ mice were generated on the *Thy1*^YFP/YFP^ background. Two age-matched cohorts of mice were generated – 1) 40 *Thy1*^YFP/YFP^::*CR3*^+/+^ and 40 *Thy1*^*YFP/YFP*^::*CR3*^-/-^ (20 male, 20 female), and 2) 40 *Thy1*^YFP/YFP^::*CR3*^+/+^ (male only). Female animals were not selected for estrous cycle synchronization.

### LA1 injections

Animal cohort 2, consisting of 40 *Thy1*^YFP/YFP^::*CR3*^+/+^ male mice, were randomly assigned to either LA1 (Leukadherin-1; S8306, Selleckchem, TX) or) or vehicle (6% DMSO (Dimethyl Sulfoxide, ACS; 191418, MP Biomedicals, PA) in sterile saline with 1% Tween-20 (BP337500; Fisher Scientific, NJ)). Injected animals were subjected to the same experimental timeline and behavior as cohort 1 and tissue from cohort 2 was processed and stained together with cohort 1. All further methods describe both cohort 1 and 2’s treatment. For further LA1 injection details see ‘Supplementary Materials’.

### Irradiation

Cranial irradiation procedures were implemented as previously described (eight-week old, male and female mice, and 10 Gy head-only γ-irradiation) (36). Prior to radiation, all animals were randomly assigned to sham or radiation groups, given a numerical code, and the experimenter was blinded to all subsequent analyses.

### Behavioral Tasks

Fourteen days prior to testing, animals were moved to a reverse 12:12 h light:dark cycle to allow behavioral testing in the day-time during the animals awake (dark) period. Three days prior to testing, animals were brought from the housing room to the testing room and acclimated to handling each day. Ten animals per group were used for all behavioral tasks. All apparatus surfaces and objects were cleaned between each animal with 75% ethanol. Any-maze behavioral tracking software (v6.1, Stoelting Co., IL) was used to record all animal videos and automatically scored open field and fear conditioning tasks while Lashley III maze and novel object recognition were scored manually. Behavioral tasks consisted of open field (OF), novel object recognition (NOR), Lashley III maze (LIII), and contextual fear conditioning (FC) with two days of extinction. For detailed behavioral task explanations and schematics see ‘Supplementary Materials’.

### Tissue Preparation

Animal brain tissue was processed as previously described (36) with the exception that mice were sacrificed 45 d post-irradiation after completion of behavioral tasks.

### Immunohistochemistry

Five animals were selected at random from each group; cohort 1) 20 *Thy1*^YFP/YFP^::*CR3*^+/+^ and 20 *Thy1*^*YFP/YFP*^::*CR3*^-/-^ (male / female, sham / irradiated) and cohort 2) 20 *Thy1*^YFP/YFP^::*CR3*^+/+^ (male; DMSO / LA1, sham /irradiated). Brain sections were processed as previously described (36) using primary antibodies Iba1 (Rabbit; Wako 019-19741; 1:2000), CD68 (Rat; Bio-Rad MCA1957; 1:500), CD11b (Rat; Bio-Rad MCA74G; 1:500) and secondary antibodies Alexa-Fluor Goat x Rabbit 647 (Invitrogen, A21245) and Goat x Rat 594 (Invitrogen, A11007) diluted at 1:2000.

### Confocal Imaging

Following immunostaining, coronal sections containing the anterior portion of the dorsal hippocampus (−2.0 to −2.5 mm from Bregma) were imaged using a Nikon A1R HD Laser Scanning Confocal Microscope (Nikon, Tokyo, Japan). Using Thy1-YFP fluorescence as a reference and framing, three z-stacks per animal were captured across the overlying molecular layer of the dentate gyrus with comparable sections selected across animals. Each image contained the stratum lacunosum moleculare as a top boundary and dentate gyrus granule cell layer as a bottom boundary. For detailed confocal parameters see ‘Supplementary Materials’.

### Imaris Volumetric Reconstruction

The brain is a complex 3D space and information is lost upon compressing the z-stack into a max projection. In order to address this issue, we imported confocal images into Imaris (v9.5; Bitplane, Belfast, U.K.) and performed volumetric analyses on structures in 3D space. A region of interest (same dimensions across all channels) was applied to Iba1, CD68, and CD11b images to mask the granule cell layer and limit analysis to the molecular layer. For detailed Imaris methods see ‘Supplementary Materials’.

### Dendritic spine quantification

This analysis was replicated from previously published methods (36,37). Thy1-YFP confocal images were analyzed using a minimum length of 40 μm per dendrite with a set magnification of 25 pixels per μm (refers to RECONSTRUCT’s magnification setting). Across five z-stacks, a total dendritic length of 400 μm was analyzed per animal with a total of five animals per sex per group. After counting individual spines, spine density was calculated per animal by dividing the total spine number by the total distance analyzed across images.

### Statistical analyses

Data are presented as mean ± SEM with all statistical analyses carried out in Graphpad Prism 8.1.2 (GraphPad Software, CA) to evaluate differences between sham-irradiated and irradiated, genotype, sex, and injectate differences. Data were analyzed by two-way ANOVA (Sholl analysis, CD11b, and LA1 data) or three-way ANOVA (all other analyses) followed by a *post hoc* Holm-Sidak multiple comparisons correction test. Data were tested for normality using the Shapiro-Wilk test and passed with a p > 0.05. P = 0.05 was considered significant (* p < 0.05, ** p < 0.01, *** p < 0.001, **** p < 0.0001).

## Results

### Irradiation-associated behavioral deficits in male WT mice are not apparent in female WT or CR3 KO animals

To investigate the effects of radiation on cognitive function, mice were subjected to a battery of behavioral tasks (Fig. 1-3, Supplemental Fig. 1-3). While WT mice did not display any differences in field preference or distance traveled following irradiation in the OF, KO mice displayed a reduction in peripheral preference and locomotor activity suggesting a lower anxiety or activity level (Supplemental Fig. 1). NOR, thought to test hippocampal-dependent memory and a rodent’s natural inquisitiveness (38), indicated a significant radiation-mediated decrease in male WT discrimination (Fig. 1b – p < 0.0001). This irradiation effect was not displayed in female mice and was absent in male KO mice (Fig. 1b – p = 0.001) suggesting that radiation-associated object discrimination impairment is limited to male WT mice.

**Figure 1.**
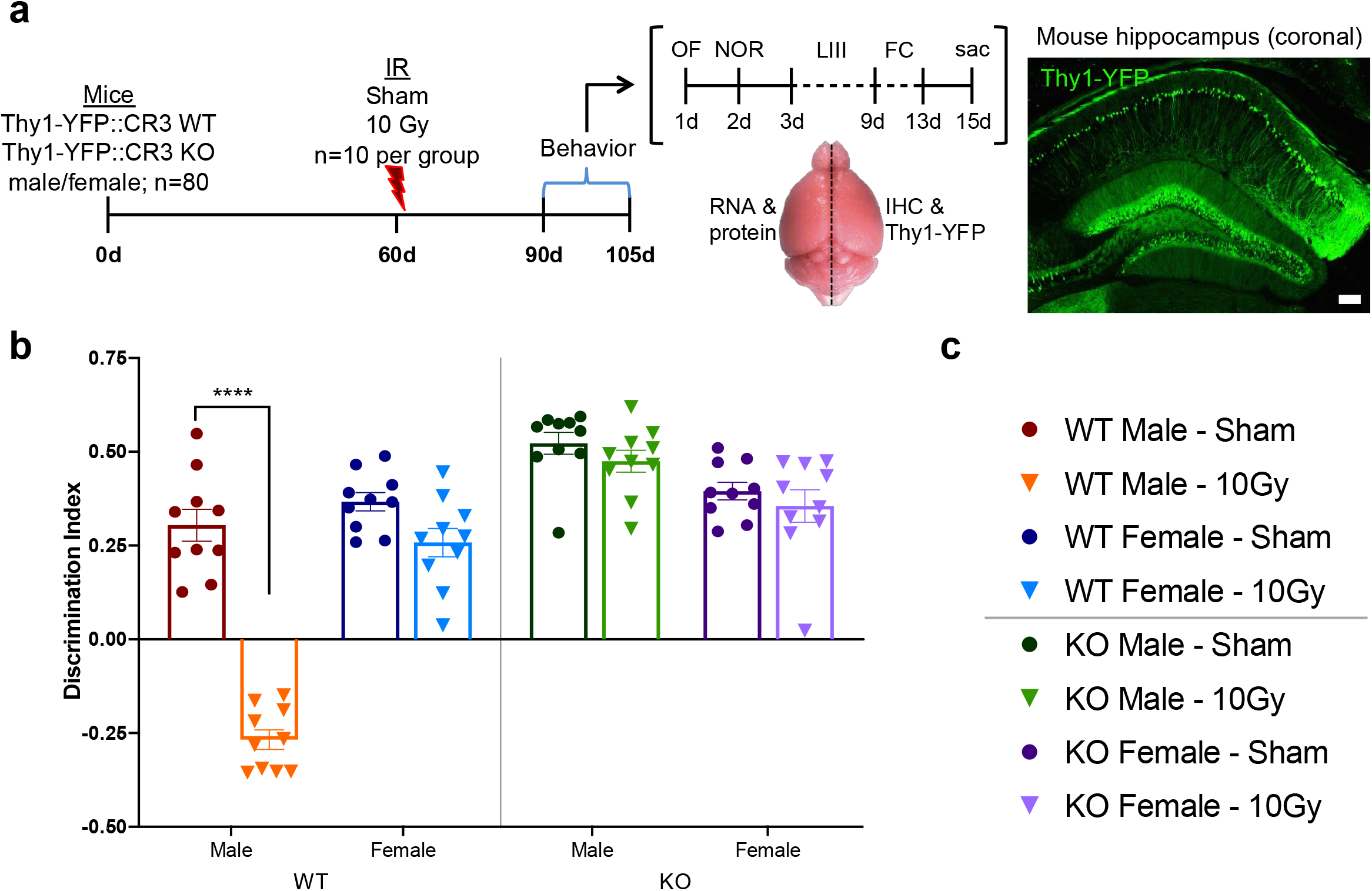
Experimental Paradigm and Novel Object Recognition (NOR) tasks. **a**) Basic schematic of experimental timeline and task workflow. **b**) Discrimination Index for NOR demonstrated a significant irradiation-mediated deficit in male WT mice that was absent in the other groups. **c**) Legend displaying color, symbol, and experimental groups that will remain consistent in Figures 2-5. n = 10 per group; **b**) three-way ANOVA followed by multiple comparisons correction, * p < 0.05, **** p < 0.0001.

Next, the LIII maze was utilized to test hippocampal-dependent memory and route learning over a period of 7 daily trials (39). Male WT sham mice performed the task with a 90% completion rate while only 20% of irradiated male WTs completed the task. Further, female and KO groups did not demonstrate an irradiation response; however, completion rates were 40% and lower suggesting male WT sham mice were more adept at this particular route-learning task (comparison of “survival” curves, log-rank; Fig. 2a, b – Mantel-Cox test: p = 0.007, test for trend: p = 0.0094). Notably, the number of errors between groups was consistent across trials suggesting that all animals learned comparably, but male WT sham mice learned more quickly and made fewer mistakes (Fig. 2c, d).

**Figure 2.**
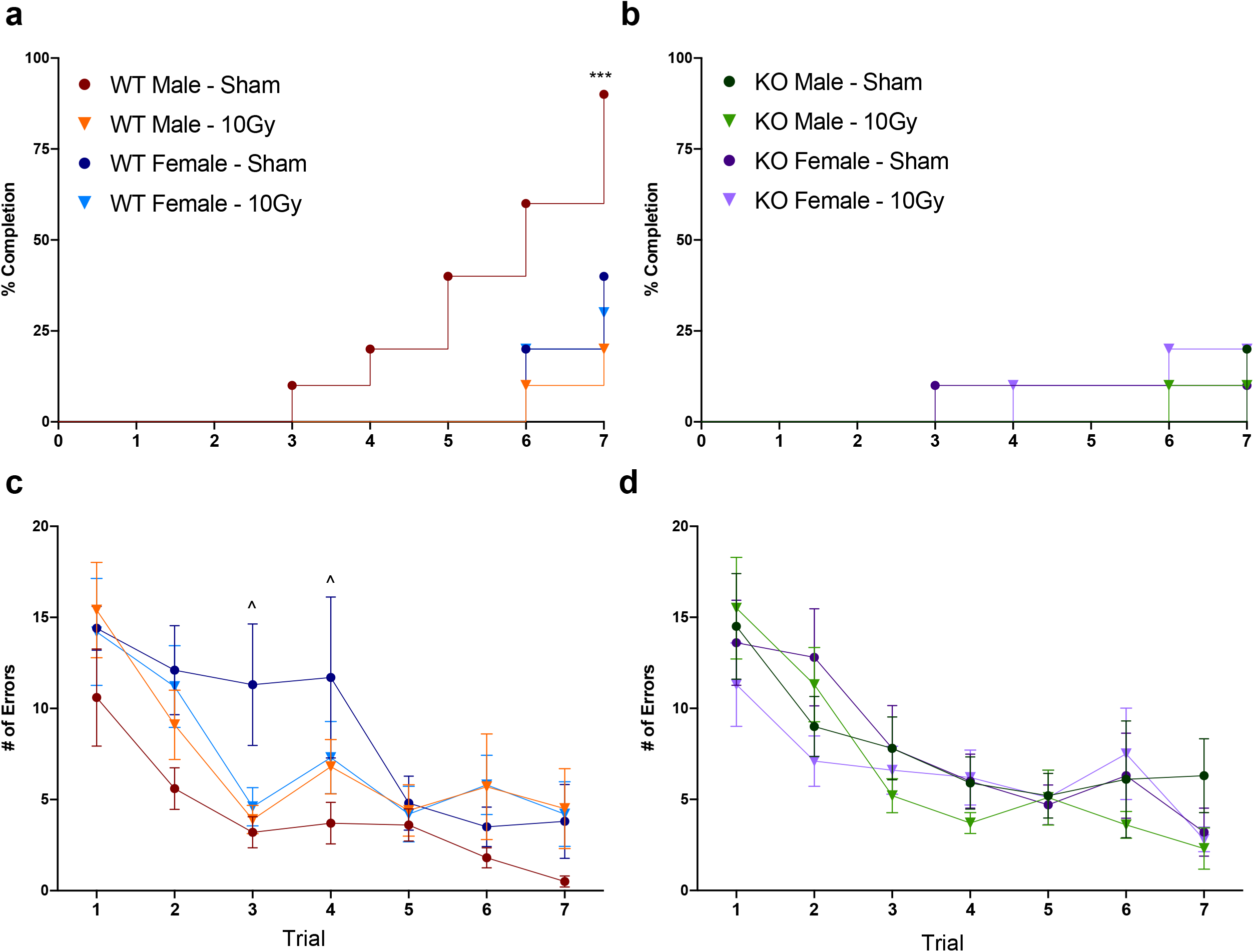
Lashley III Maze (LIII). **a, b**) Percent completion is plotted as a reverse ‘survival’ curve across the daily trials and shows that male WT sham mice completed the task efficiently while the other groups did not. Although many of the animals did not meet completion criteria, the number of errors per trial was similar across groups (**c, d**). n = 10 per group; **a, b**) Mantel-Cox test and test for trend; **c, d**) three-way ANOVA followed by multiple comparisons correction, */^ p < 0.05, *** p < 0.001. Sham vs irradiated significance: * male, ^ female.

The last task, context fear conditioning with extinction, tests the animal’s ability to learn a shock-tone pair and then dissociate the adverse conditioning pair, measured by freezing response, over an extinction period (40). During the Context test, baseline percent freezing was lower in KO mice when compared to WT mice (three-way ANOVA; Fig. 3a, b – F (1, 72) = 50.20, p < 0.0001); however, during the Tone test, there were no significant differences in freezing response with or without the tone. Day 1 of Extinction demonstrated a radiation effect on the ability of WT mice to unlearn the fear response that was not present in the KO animals (three-way ANOVA; Fig. 3a, b – IR: F (1, 66) = 6.972, p = 0.0103; genotype: F (1, 66) = 0.3561, p = 0.5527). Similarly, day 2 of Extinction indicated that irradiated WT animals fail to forget the associated response but female mice demonstrated a non-significant trend to decrease percent freezing in trials 3-5. KO animals could neither forget nor were affected by radiation and showed a similar percent freezing to Extinction day 1 (three-way ANOVA; Fig. 3a, b – IR: F (1, 334) = 34.12, p < 0.0001; genotype: F (1, 334) = 56.55, p < 0.0001).

**Figure 3.**
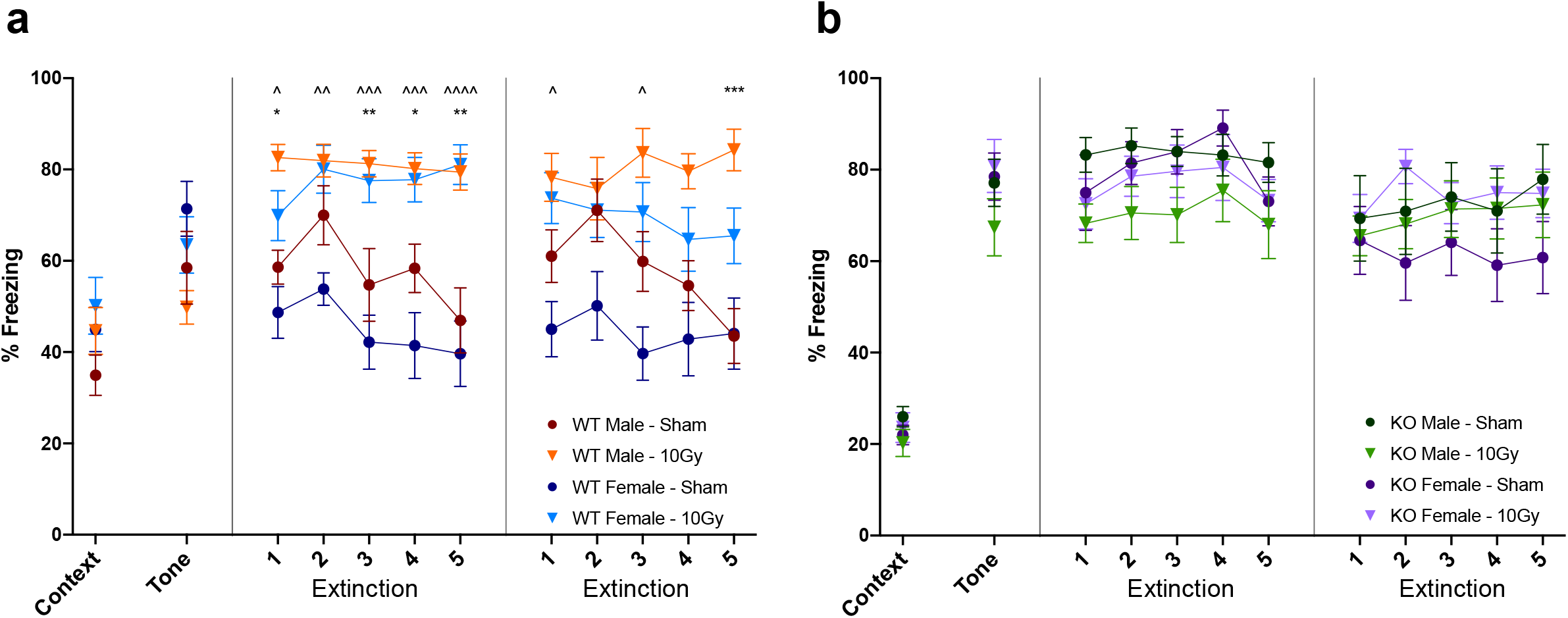
Context Fear Conditioning (FC) with extinction. Each phase is measured by a freezing percentage and plotted as conditioning, context, tone test, and extinction (**a, b**). **a**) Displays WT mice and demonstrates a significant irradiation-mediated effect in extinction while **b**) KO animals did not show an irradiation effect or extinction. n = 10 per group; **a, b**) three-way ANOVA followed by multiple comparisons correction, ^/* p < 0.05, ^^/** p < 0.01, ^^^/*** p < 0.001, ^^^^ p < 0.0001. Sham vs irradiated significance: * male, ^ female.

### Dendritic spine density is significantly decreased in only male WT mice following irradiation

To further investigate irradiation-mediated effects, Thy1-YFP positive spines were manually counted to measure spine density (Fig. 4). Male WT spine density was significantly reduced after radiation exposure (Fig. 4d – p < 0.0001). In contrast, male KO mice showed a significant increase in basal spine density when compared to WT males (Fig. 4d – p < 0.0001) and no irradiation-associated change in spine density (Fig. 4d – p = 0.879). Female mice, both WT and KO, did not demonstrate irradiation-mediated spine density loss, indicating a lack of spine susceptibility in female mice (Fig. 4d; p > 0.05). Interestingly, when comparing WT sexes, male mice demonstrated a lower baseline spine density than female mice that was not apparent in KO mice (Fig. 4d – WT: p = 0.0001; CR3 KO: p = 0.454). These data indicate that CR3 plays a role in irradiation-mediated spine loss in male mice in the molecular layer of the dentate gyrus.

**Figure 4.**
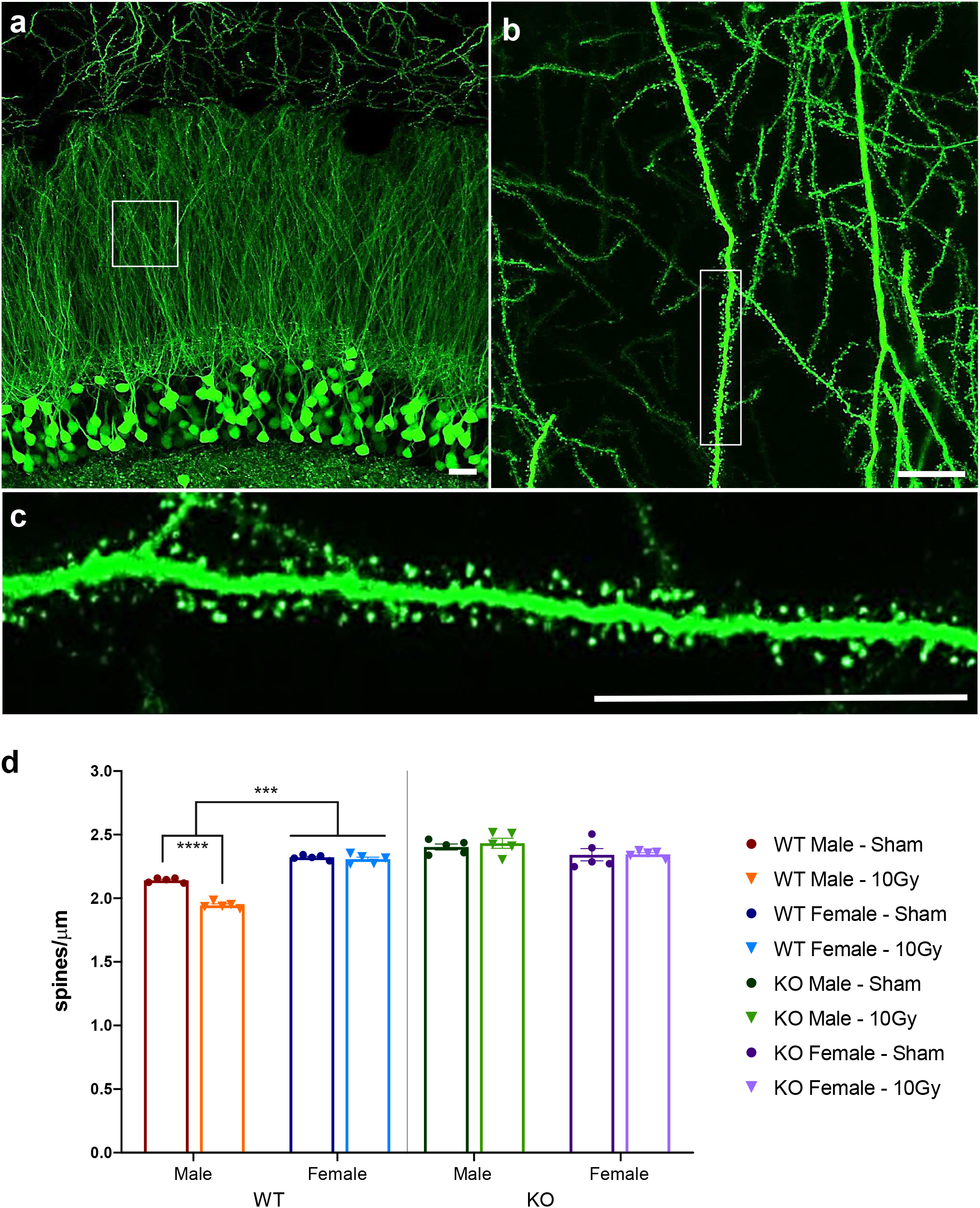
Representative Thy1-YFP images and quantification of spine densities. Representative sham-irradiated Thy1-YFP male confocal z-stack displaying **a**) the hippocampal molecular layer and dentate gyrus granule cells with **b**) magnified dendrite inset and **c**) further magnification showing individual spines. **d**) Quantification of spine density across all groups demonstrated a significant irradiation-mediated loss of spines in male WT mice but not female or KO mice. **d**) Three-way ANOVA followed by multiple comparisons correction, *** p = 0.0001, **** p < 0.0001. Scale bar: **a, b, c**) 10 µm.

### Irradiation-mediated increases in CD68 and CD11b are sex specific

To quantify modifications in microglial activity via volumetric analysis, tissue was stained for the classical microglial markers, CD68 and CD11b (Fig. 5a). Post-irradiation, CD68 and CD11b volume was significantly increased in male mice but not female mice (Fig. 5b, CD68 – male: WT, p < 0.0001; KO, p < 0.0001; female: WT, p = 0.0614; Fig. 5c, CD11b – male: WT, p = 0.0032). While irradiation provoked a slight, non-significant CD68 increase in female WT mice, it was substantially less than the male response and both CD68 and CD11b baseline levels were lower than male WT counterparts (sham WT; Fig. 5b – CD68, p = 0.0164; Fig. 5c – CD11b, p = 0.0032). We also quantified microglial morphology using an Imaris 3D Sholl analysis and demonstrated a modest radiation effect (Supplemental Fig. 4a-d).

**Figure 5.**
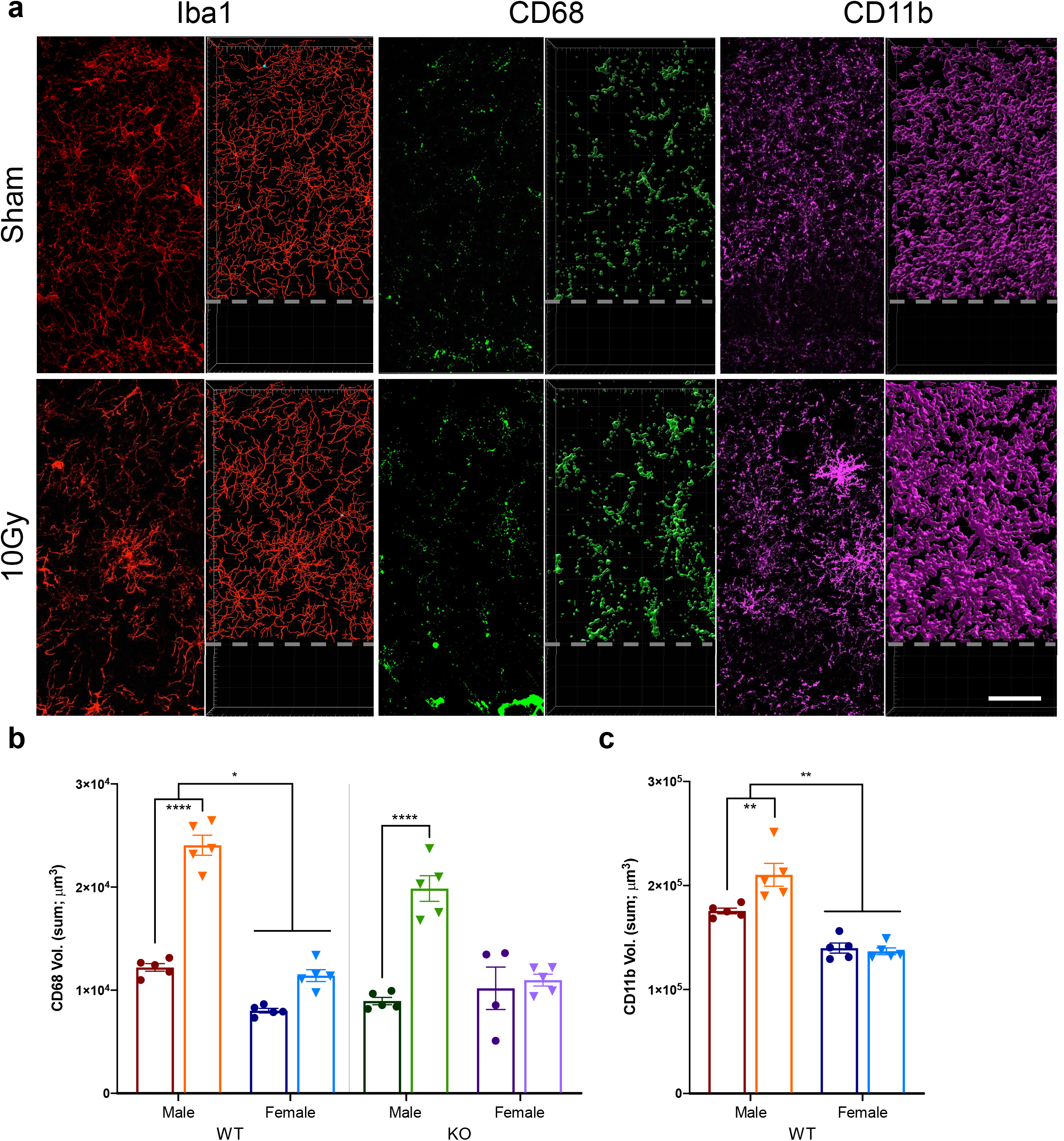
Representative confocal images of microglial markers. Iba1, CD68, and CD11b with right half panel displaying Imaris volumetric reconstruction in male sham and irradiated mice (**a**). Quantification of **b**) CD68 and **c**) CD11b volume demonstrated a significant irradiation-mediated increase in male mice but not female mice. n = 5 per group; **b, c**) three-way ANOVA followed by multiple comparisons correction, * p < 0.05, ** p < 0.01, **** p < 0.0001. Scale bar: **a**) 50 µm. Dotted grey line denotes uniform region of interest set across images to remove the dentate gyrus granule cell layer.

### LA1 treatment ameliorates irradiation-mediated deficits in novel object recognition and LIII completion rate

FC behavioral tasks (Fig. 6e) did not reveal an irradiation response in vehicle and LA1 treated mice. However, an irradiation-mediated deficit was demonstrated in NOR (Fig. 6c – vehicle: p < 0.0001; IR: p < 0.0001) and LIII criteria completion (Fig. 6d – Mantel-Cox test: p = 0.0114, test for trend: p =0.0083) in vehicle treated mice, which was ameliorated by LA1 treatment. Sham animals and LA1 irradiated animals demonstrated an LIII completion rate of 60% while only 20% of irradiated vehicle mice completed the task.

**Figure 6.**
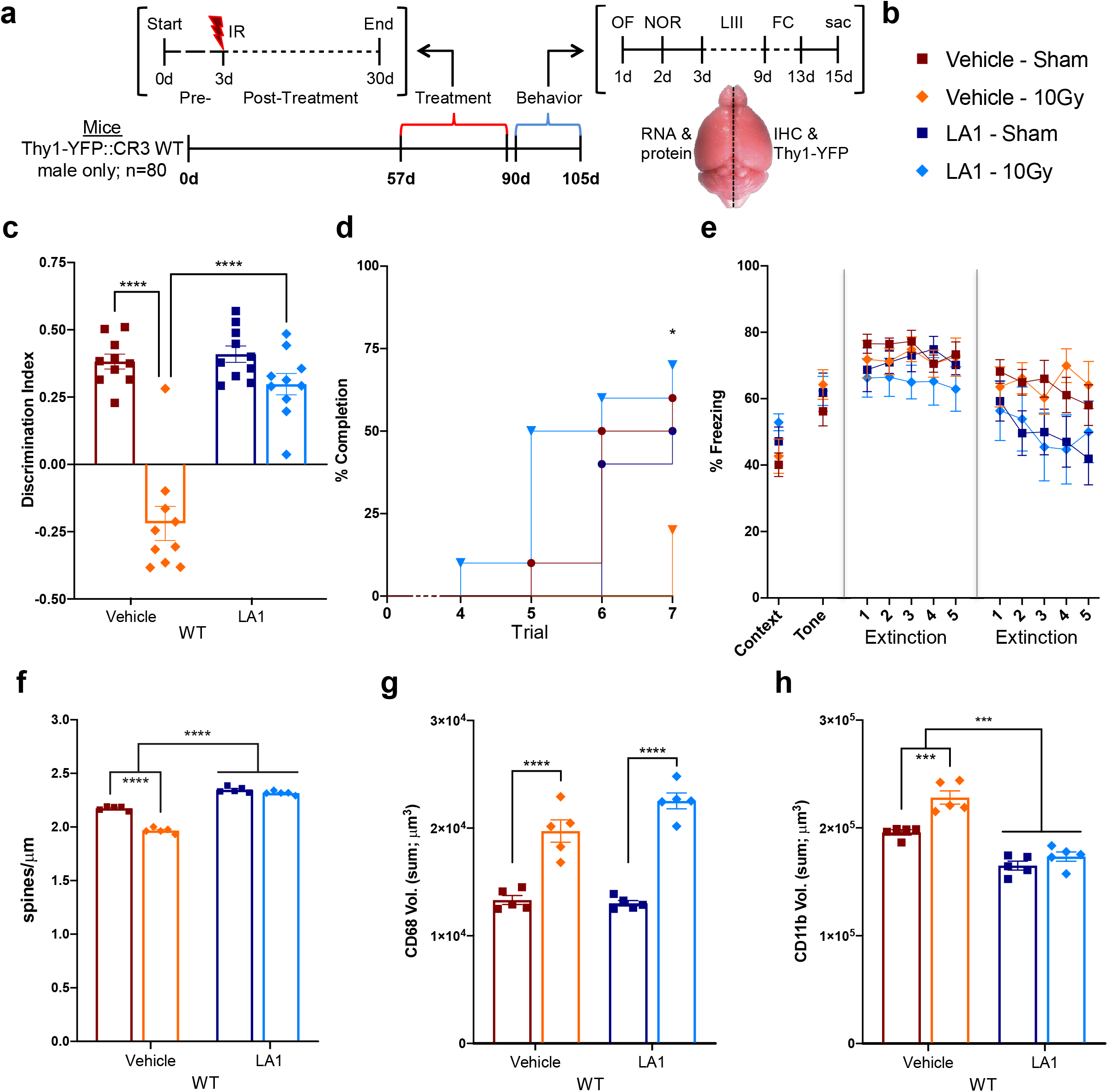
Experimental timeline for LA1 and vehicle treated male mice. **a)** Experimental paradigm identical to the previous experiment with the exception of daily LA1 injections for 30 d. **b**)Modified legend with new symbols for this cohort of mice. NOR (**c**) and LIII (**d**) demonstrated a significant irradiation-induced deficit in vehicle task performance that was prevented in LA1 treated mice while **e**) FC did not demonstrate a difference in between groups. Quantification of confocal images displaying **f**) Thy1-YFP spine density, **g**) CD68, and **h**) CD11b. n = 10 (**c-e**) and n = 5 (**f-h**) per group; **c, e-h**) two-way ANOVA followed by multiple comparisons correction; **d**) Mantel-Cox test and test for trend; * p < 0.05, *** p < 0.001, **** p < 0.0001.

### Irradiation-mediated dendritic spine loss is prevented in LA1 treated mice

Thy1-YFP dendritic spine density was significantly reduced in vehicle treated mice following radiation exposure (Fig. 6f – p < 0.0001) while LA1 treated animals did not demonstrate a radiation effect (Fig. 6f – p > 0.05). Additionally, LA1 treatment was associated with increased basal spine density when compared to vehicle controls (Fig. 6f – p < 0.0001). These data indicate that LA1 treatment plays a role in modulating irradiation-mediated hippocampal spine loss in male mice.

### LA1 effects on microglia

Microglial CD68 volume was significantly increased in vehicle and LA1 treated mice following radiation (Fig. 6g – p = 0.0001). CD11b volume was upregulated in vehicle (Fig. 6h – p = 0.0004), but this increase was prevented in LA1 treated mice (Fig. 6h – p = 0.206) and LA1 treated mice demonstrated a baseline decrease in CD11b volume when compared to vehicle treated mice (Fig. 6h – p = 0.0005). Additionally, microglial 3D Sholl analysis demonstrated a modest radiation and LA1 treatment effect (Supplemental Fig. 4e-g).

## Discussion

### Behavioral tasks reveal irradiation-mediated cognitive deficits in male WT mice but no apparent deficits in female WT or CR3 KO mice

The selected behavioral tasks were chosen to limit aversive stimuli and, with the exception of fear conditioning (run last due to aversive foot shock), all tasks maintained low stress conditions. In the NOR task reactivity to a novel object decreases with age, injury, or irradiation and likely results from damage to the perirhinal cortex and hippocampus which manifests as memory impairment (38). We found an irradiation-mediated deficit in novel object recognition, but only in male WT mice (Fig. 1b). In contrast, female WT mice and KO mice of both sexes did not display an associated radiation effect and preferred the novel object over the familiar one. It is important to note that others have failed to find deficits or demonstrated enhanced task performance (41) following radiation; these discrepancies are likely a result of differences in radiation dose, sex, age, genotype, and general task performance.

Many rodent behavioral tasks designed to test spatial learning and route memory rely on visual cues and aversive stimuli to motivate task completion. The use of these motivators may be confounding as age- and strain-dependent visual acuity and stress/pain sensitivity are highly variable (39). To minimize these effects, the Lashley III maze was utilized as a low-stress, hippocampal-dependent navigational task with a pseudo-home cage as a motivational reward (Supplemental Fig. 2). This neutral motivation avoids further confounds such as food or water deprivation, food reward seeking, odor cues, or experimenter interference. Similar to a previous study (39), nearly all male WT sham mice (90%) were able to complete the maze in 7 days (defined as running the maze with 0 or 1 error on two consecutive daily trials).; however, only 20% of male WT mice completed the task post-irradiation, reflecting an effect of radiation on navigation. The other groups, female WT and KO mice, displayed an overall reduction in ability to complete the maze, with or without irradiation (< 40%; Fig. 2a, b). It is important to note that all animals had similar trends in the number of errors and had fewer than 7 errors by day 7, indicating that all animals were able to learn the task (Fig. 2c, d). Together, these data suggest that male WT mice are more proficient (task completion) at route learning than female WT mice and sensitive to irradiation-mediated cognitive deficits. Because female WT mice and KO mice of both sexes performed poorly on the LIII task, we were not able to detect an irradiation-mediated effect in these groups. Further studies are needed to determine whether this was due to an inability to learn the task, enhanced radio-resistance, or a need for additional days to learn (39).

The last behavioral task, contextual FC with extinction, has multiple phases that test hippocampus, pre-frontal cortex (PFC), and amygdala pathways in the acquisition, recollection, and dissociation of a paired conditioned (tone)-unconditioned (foot shock) stimulus (Fig.3; Supplemental Fig. 3). The first stage, Context, is hippocampal-dependent and tests the extent to which an animal can associate the encoded context (environment) and aversive foot shock without the shock or tone cue. The second Tone stage is context-independent (novel environment) and tests non-hippocampal pathways in an animal’s recollection of the shock when presented with the tone (likely amygdala- and PFC-dependent). The third and fourth stages, Extinction day 1 and 2, test hippocampal-dependent spatial context retrieval of the shock memory prompted by the tone, and after repeated presentation of the tone, measures the extinction or dissociation of the fear response (40). Our results demonstrate that WT mice did not exhibit an irradiation- or sex-mediated difference in Context and Tone stages; however, after repeated exposure, irradiated WT mice failed to dissociate the fear response and freeze relative to sham WT animals by day 2 of Extinction (Fig. 3a). In contrast, CR3 KO mice failed to extinct and did not display an irradiation-mediated response (Fig. 3b). When compared to WT mice, KO mice demonstrated: a) a significantly lower Context freezing level, indicating a reduction in context-dependent fear learning; b) a higher Tone freezing response, suggesting an increase in context-independent conditioning; and, c) a higher Extinction freezing percentage, similar to irradiated WT mice, indicating an inability to dissociate the fear response after repeated presentations of the tone. Notably, female WT mice start the extinction task close to Context freezing levels (i.e. already extinct), suggesting reduced contextual retrieval after 48 h or greater ability to dissociate the shock-tone response (Fig. 3a). Similar to NOR, these fear conditioning results agree with some studies (42), but conflict with others (14), and are likely due to the differences in animal handling and protocols as previously mentioned.

### Cranial irradiation leads to a significant loss in spine density in male WT mice that is prevented in female WT and CR3 KO mice

The ability to structurally modify and remodel dendritic spines affords flexibility in connectivity and permits allocation of experience-dependent memories (8). While direct translation of spine number to circuit functionality and memory formation is poorly understood, it is reasonable to speculate that dendritic spine loss can lead to cognitive impairment. In this study, our data demonstrate a significant reduction in spine density in male WT mice (−9.2%) following radiation that was prevented in male KO mice and absent in female mice (Fig. 4d). These results corroborate our prior findings (36) that dendritic spine loss is sex-dependent and CR3 deficiency prevents the loss following irradiation. NOR data suggests that spine loss correlates with a cognitive deficit in male WT mice that is ameliorated in male CR3 KO mice. This is further supported in female WT mice where there is no dendritic spine loss following irradiation and cognitive impairment does not manifest. Unfortunately, we were not able to verify similar protection in females using our LIII route-learning task due to poor behavioral task performance. In addition, while our results agree with previous studies performed in male mice 30 days post-irradiation (10 Gy) (10,11,36) the severity of spine loss is less pronounced in this current study. This may be due to different staining methods and quantification analyses implemented across the studies – Golgi-Cox stain with manual counting (10,36), Thy1-YFP positive spines with manual counting (current study), and Thy1-YFP positive spines with Imaris volumetric reconstruction and automatic counting (11).

### Irradiation induces elevation in microglial activation markers in male, but not female mice

Following cranial irradiation, DNA damage and cellular injury provoke a shift in inflammatory signaling and microglial reactivity. As resident CNS macrophages, a classic response to injury and inflammation is increased microglial phagocytosis and debris clearance (17,20). Two markers involved in phagocytosis and activation, CD68 and CD11b, were significantly upregulated in response to radiation in male WT, but not female WT mice. This suggests that irradiated male WT mice have an increased potential for CD68- and CR3-mediated phagocytosis of opsonized material when compared to female WT mice (Fig. 5b). A similar CD68 elevation occurred in male KO mice, indicating comparable phagocytic capacity; however, KO mice are deficient in CD11b and therefore lack CR3-mediated recognition and removal of opsonized material (Fig. 5c). The observed sex differences may be due to the more pronounced immune capacity and inflammatory phenotype when comparing male to female mice (43).

Consistent with our results, studies using high-LET particle irradiation demonstrated sex differences in microglial activation, synaptic modifications, and cognitive deficits (44,45), with females being relatively protected. While findings regarding sex differences in rodents support an association between microglial activation, synapse loss and cognitive deficits, they may not reflect responses in patients, as several clinical studies indicate greater sensitivity in females, at least for childhood survivors of clinical radiotherapy (46).

### Leukadherin-1 (LA1) treatment prevents irradiation-mediated cognitive deficits and spine density loss

To complement the use of a global CR3 KO model and avoid the potential influence of CR3 deficiency on neuron-glia signaling and interaction in development and aging, male mice were injected with vehicle or the small molecule, CD11b agonist LA1 (47) to determine the role of transient CD11b dysfunction in irradiation response. LA1 is thought to bind the αA-domain of CD11b and lock it in an active conformation or intermediate affinity that leads to endocytosis without initiating intracellular signaling; e.g., signaling that would result in myeloid trafficking and recruitment, phagocytosis, and cytokine production (48,49). Additionally, LA1 readily crosses the BBB to target microglial CR3 (50), and synergizes with radiation to suppress breast tumor growth (*in vitro*) and enhance survival in mouse models of human cancer by slowing tumor progression (47).

Following irradiation, vehicle and LA1 treated animals exhibited no difference in OF preference or distance traveled; indicating LA1 does not induce abnormal, anxiety-like behavior when compared to vehicle controls (Supplemental Fig. 1f, g). In the NOR task, vehicle treated male mice demonstrated significant irradiation-mediated deficits in novel object recognition that were absent in LA1 treated animals (Fig. 6c). Similarly, in the LIII maze, there was a significant irradiation-mediated effect in vehicle treated mice that was prevented in LA1 treated mice (Fig. 6d). Unfortunately, it is difficult to further extrapolate on the context fear conditioning data, as both vehicle and LA1 treated mice performed similarly and did not display an irradiation-mediated effect in Extinction trials. This could be due to a lack of conditioning and poor task performance or could indicate that the vehicle treatment alone afforded some benefit.

Consistent with behavioral findings, we demonstrated decreased spine density in vehicle treated mice post-irradiation that was prevented in LA1 treated mice (Fig. 6f). Additionally, irradiation-mediated spine loss was associated with increased CD68 (Fig. 6g) while LA1 prevented the irradiation-induced increase of CD11b displayed in vehicle treated animals (Fig. 6h). These data suggest that while phagocytic capacity is elevated post-irradiation in either treatment, CD11b upregulation is prevented by LA1 and may protect against spine loss and associated cognitive dysfunction. Interestingly, basal spine density levels were significantly increased in LA1 treated mice when compared to vehicle suggesting effects on normal spine turnover. It is important to note that although LA1 crosses the BBB and targets microglial CD11b, it also affects peripheral cells. Thus, we cannot fully differentiate microglial-specific effects from effects on peripheral immune cells.

In this second experiment utilizing LA1, the data corroborate the results from the first experiment: irradiation-mediated dendritic spine loss is driven by elevated microglial CR3-dependent phagocytosis and is associated with cognitive impairments in male WT mice. The second experiment also identifies LA1 as a potential therapeutic approach that prevents irradiation-mediated dendritic spine loss, CD11b elevation, and associated hippocampal-dependent spatial memory deficits.

## Conclusion

Cranial radiation therapy is the leading mode of therapeutic treatment for primary and metastatic malignancies in the brain. Unfortunately, cranial radiation is associated with significant late-delayed, progressive, and irreversible cognitive deficits that are currently untreatable and permanently reduce patient quality of life. Due to the detrimental neurocognitive complications and the critical role of synapses in the efficient function of neuronal circuits, our study focused on the dendritic spines in the hippocampus, a region critical for learning and memory. In this study, we corroborate previous findings (36) and provide further insight into the mechanisms connecting irradiation-mediated cognitive decline, spine loss, microglia, and the complement pathway. Additionally, we identify a potential therapeutic, LA1, which may have direct translational benefit to cancer patients by reducing irradiation-induced tissue damage and improving cognitive outcomes. Because similar structural changes and cognitive deficits are observed following heavy ion or proton radiation (12,13,16), this therapeutic approach may also be beneficial to patients undergoing particle radiation therapy as well as astronauts during deep space travel. Clearly, additional experiments exploring more clinically relevant radiation dosing paradigms (e.g. fractionation) and the persistence of benefits need to be performed. Future studies testing the effects of inhibiting complement components on tumor killing are also required.

## Acknowledgements

We thank Lee Trojanczyk, Laura Owlett, Dawling Dionisio-Santos, Berke Karaahmet, and Ania Majewska for input and discussion throughout these experiments. We also thank the Director of the Confocal and Conventional Microscopy Core (CCMR; University of Rochester Medical Center), V. Kaye Thomas, for use of the confocal and image acquisition discussions.

