## Supplementary Methods and Figures for "Pharmacologic manipulation of complement receptor 3 prevents dendritic spine loss and cognitive impairment after acute cranial radiation"

**Supplementary Materials**

**Leukadherin-1 Injections**

Mice were given 6 mg/kg daily intraperitoneal injections (i.p.) of LA1 or vehicle starting 3 d prior to cranial radiation (pre-treatment) for a total of 30 d. This dose was determined from previous literature (1,2) and animals displayed no overt health issues, changes in behavior, or toxicity. To prep for injections, LA1 was stored as a stock solution of 10 mg/ml in DMSO, aliquoted into tubes, and stored at -80°C for daily, single use. Fifteen minutes before injections, LA1 was removed from the freezer, thawed, and vortexed until solubilized. Five mins before injections, 360 μL of stock LA1 or DMSO was added to 5.64 mL saline/1% Tween-20, vortexed, and injected at 6 mg/kg (i.e. a 25 g mouse received 250 μL).

**Behavioral Tasks**

*Open Field and Novel Object Recognition*: The open field task provides a rapid assessment of well-defined anxiety-like behaviors since mice have a natural propensity to avoid bright, open, and unknown environments and thus preferentially explore the periphery (wall-hugging) (3). The open field task was also used as a habituation phase for the subsequent novel object task. During the OF task animals were placed in a 30.5 cm x 30.5 cm x 25.5 cm box (white floor and black walls) for 5 min. Any-maze software was set to track and score when the mouse was in the periphery verse the center of the floor as well as distance traveled. Following a 24 h period, mice were placed back into the box with two identical objects (ceramic door knobs, 6 cm in height and placed 15 cm apart). Two hours after the training phase, animals were returned to the box containing one of the previous objects (familiar) and a new object (novel) to explore for a 5 min testing phase. Novel object exploration was scored based on the time the animal spent “interested” in the object; head facing and in close proximity to the object, neck extended, vibrissae moving. Simply passing or standing on the object did not count as time spent exploring. Exploration times in the OF task were defined by a field preference where the time spent in the periphery was subtracted from the time in the center, divided by the total time – a value of 1.0 means the mouse prefers only the periphery, -1.0 is center preference only, and 0 is no preference for either space. The NOR task was defined by a discrimination index, DI: (t_1_-t_2_)/(t_1_+t_2_), where t_1_ is the novel object (NOR) time and t_2_ the familiar object (NOR) time. A DI with a positive index indicates mice prefer the novel object over the familiar one (NOR). A negative index is the opposite preference and an index of zero indicates no preference for either object.

The OF task demonstrated a significant genotype difference in which KO mice had less preference for the peripheral space and traveled a shorter distance when compared to WT mice (three-way ANOVA, genotype; Supplemental Fig. 1c – F (1, 72) = 65.40, p < 0.0001; Supplemental Fig. 1d - F (1, 72) = 49.24, p < 0.0001). Further, KO male mice demonstrated a loss of peripheral preference and decreased distance traveled when compared to female KO mice (three-way ANOVA, sex difference; Supplemental Fig. 1c – F (1, 72) = 8.518, p = 0.00471; Supplemental Fig. 1d - F (1, 72) = 13.91, p < 0.0004).

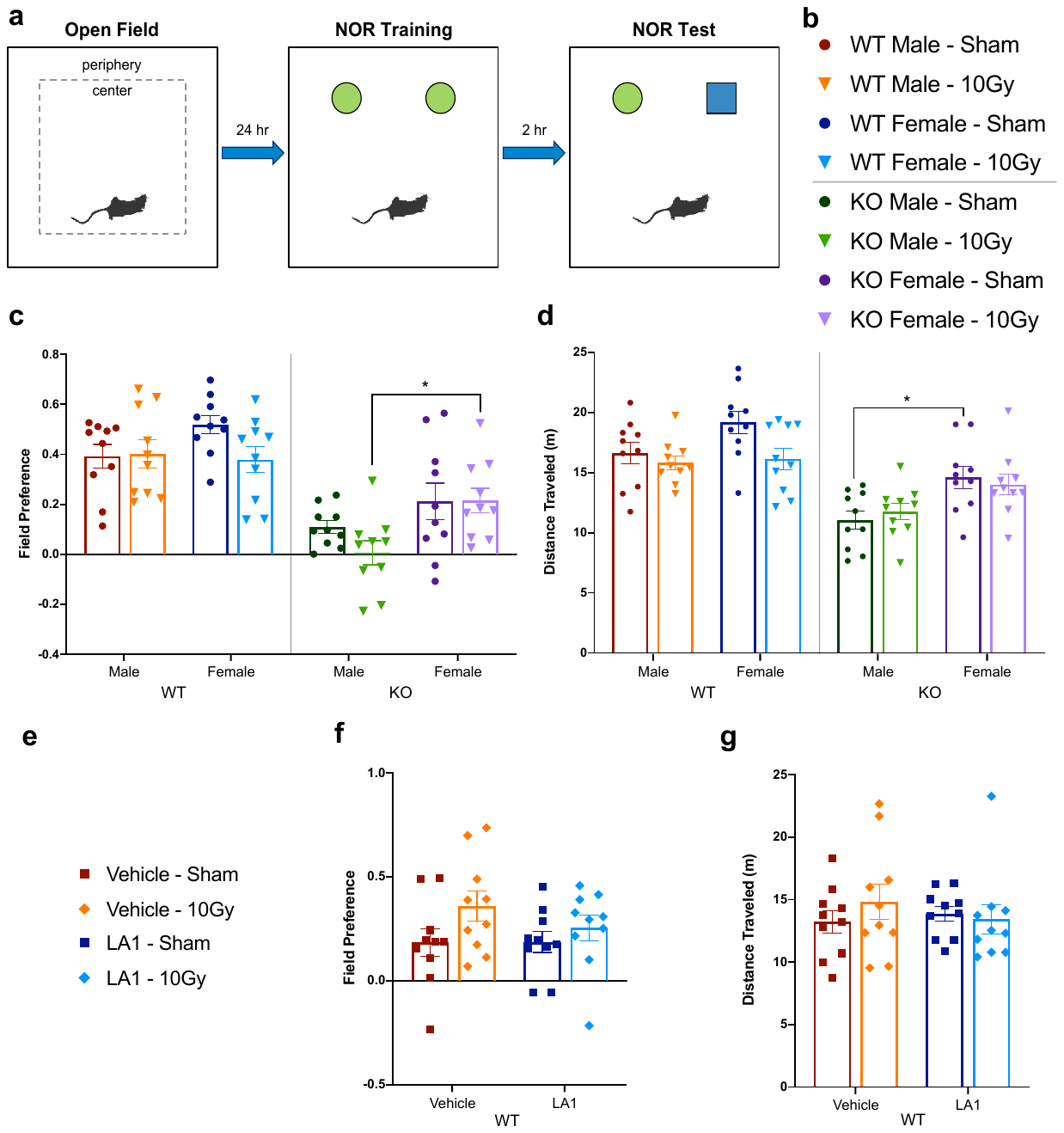

**Supplemental Figure 1. Open Field (OF) and Novel Object Recognition (NOR) tasks. a**) Basic schematic of task layout and workflow. **b**) Legend for WT and KO groups, **c**) quantification of field preference score for OF demonstrating a stronger positive score (peripheral preference) for WT mice, and **d**) total distance traveled. **e**) Legend for vehicle and LA1 groups with **f**) field preference and **g**) distance traveled demonstrating similar trends between vehicle and LA1 groups. n = 10 per group; **c, d**) three-way ANOVA and **f, g**) two-way ANOVA followed by multiple comparisons correction, * p< 0.05.

*Lashley III maze:* The apparatus, set-up, and task were replicated from a previous study (4) and implemented to assess spatial learning and route memory without the use of visual cues, food/water deprivation, or aversive stimuli. Based on the aforementioned study it took an average of 7.2 (± 1.5) trials for 2 m old C57BL/6N mice to learn the task; therefore, 7 trials, 1 trial per day, was chosen for our protocol. Due to the repeated exposure (trials) to a set route the task was considered completed when the animal performed the route with 0 or 1 error(s) on two consecutive trials. All LIII videos were manually scored due to software tracking discrepancies (low-light and shadows) so it was not possible to accurately measure distance traveled. Analysis measures consisted of criterion met (days to learn, plotted as survival) and number of errors per trial. One exception to the set-up when compared the prior study (4) was that animals were not provided with personal pseudo-home cages prior to the task; instead, a single cage with a door cut out and paper towel covering the bottom was used for each animal.

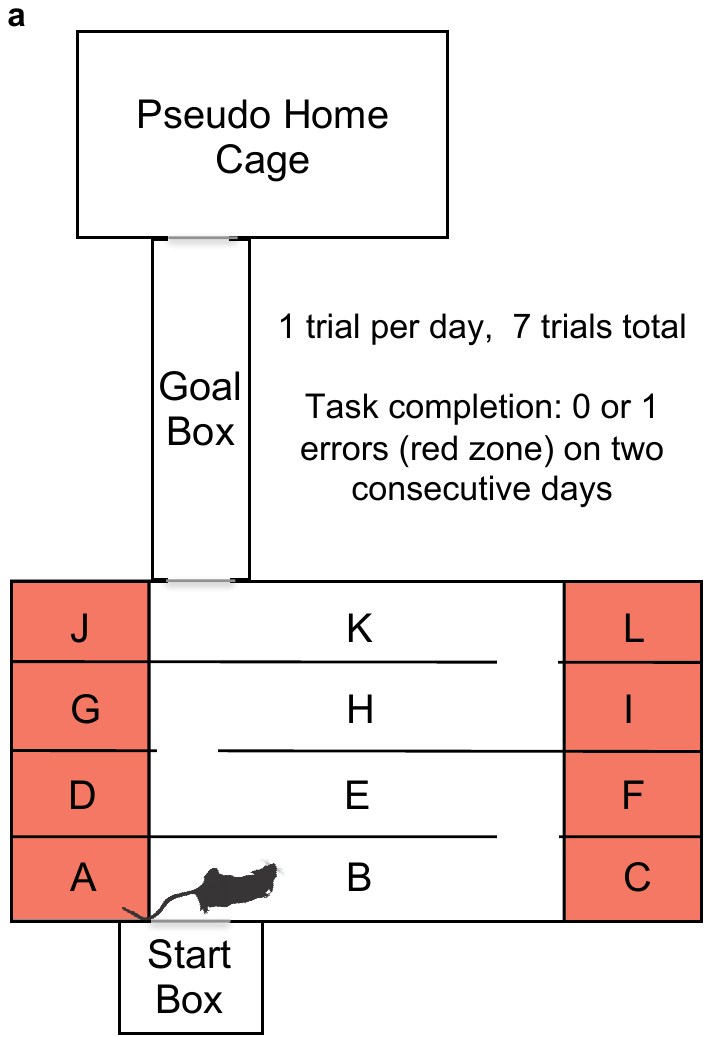

**Supplemental Figure 2. Lashley III Maze (LIII).** Basic schematic of task layout where the animal begins in the start box and explores freely through the maze. Red regions are considered ‘error’ zones as they lie outside the direct route to the Goal Box and Psuedo Home Cage. Mice are run for 7 days, 1 trial per day, and the task is considered complete when the animal performs the task with 0 or 1 error(s) on two consecutive days.

*Contextual Fear Conditioning with Extinction*: The fear-conditioning set-up consisted of a plexiglass chamber with a removable metal floor grid housed inside a large, sound-limiting box under dim, white lighting (model H10-11M; Coulbourn Instruments, PA). Details of the test are summarized in Supplemental Figure 3. On day 1, animals were placed in the context chamber for 180 s followed by 15 s of white noise (tone; 80 dB) co-terminating with a 2 s, .75 mA foot shock. This noise-shock pairing, or conditioning, was repeated for a total of 3 times with 30 s intervals in between. On day 2, 24 h later, mice were re-exposed to the context chamber and explored freely for 5 min (without tone or shock, context period). Two hours later, animals were placed in a novel context within the context chamber (a 20 cm, opaque, open-topped cylinder with clean bedding on the floor and dim, red light) for 180 s (no tone) followed by re-exposure to the tone for 180 s (tone) to test hippocampal-independent memory. Day 3 and 4 consisted of an extinction paradigm where animals were placed in the context chamber for 20 min and presented with five intermittent, 2 min long tone periods to test the ability for the animals to unlearn, or forget, the conditioned tone-shock response. All videos were recorded with Any-maze software and freezing threshold (sensitivity) values set as the recommended range (30, 40) for all animals. Analysis measures consisted of percent freezing during each of the periods from day 1 through 4.

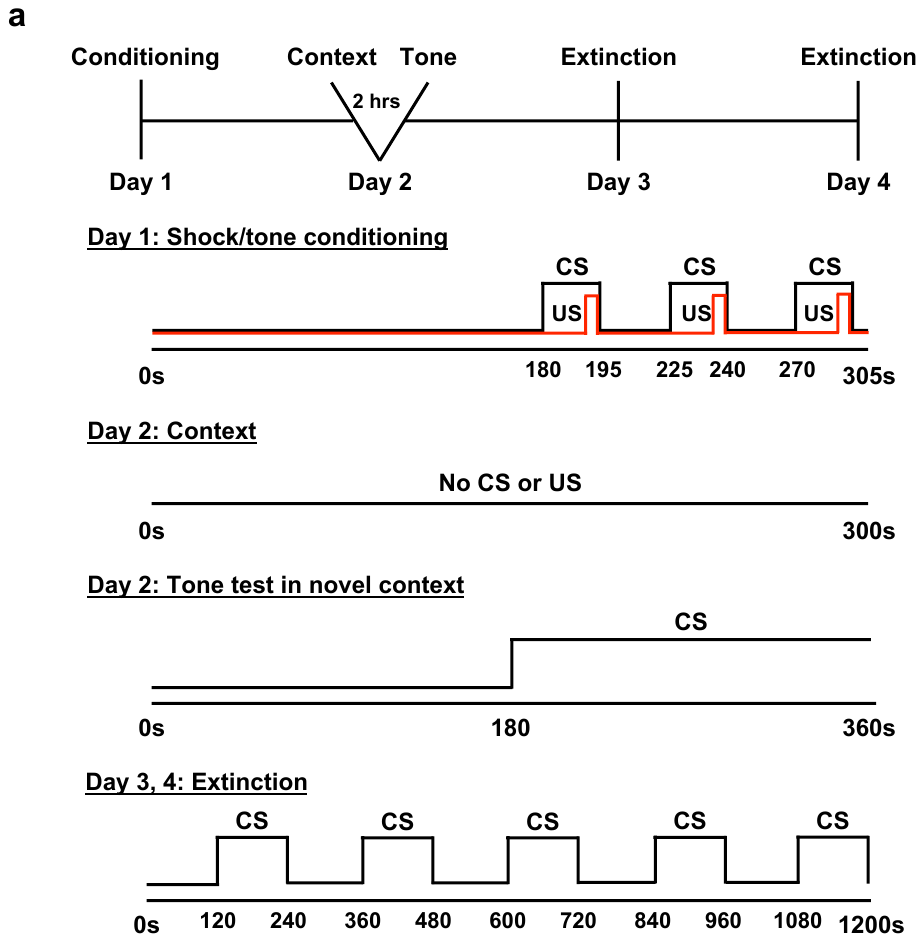

**Supplemental Figure 3.** Visual schematic of fear conditioning task.

**Confocal Imaging**

For microglial markers (Iba1, CD68, and CD11b), each z-stack was 20 μm thick (1024 x 1024 pixels) and acquired at 40x with 0.5 μm steps (0.29 μm pixel size; Nikon Apochromat Lambda LWD 40x/1,15 Water) for a total of 40 images per stack. Each image contained Iba1-647 and CD68-594 or CD11b-594 and representative images were pseudo-colored with red (Iba1), green (CD68), and magenta (CD11b) based on personal preference. For Thy1-YFP images, each z-stack was 20 μm thick (1024 x 1024 pixels) and acquired at 60x with 0.2 μm steps (0.15 μm pixel size; Nikon Apochromat TIRF 40x/1,49 Oil) for a total of 100 images per stack. Additionally, Thy1-YFP images were deconvolved (Richardson-Lucy, 20 iterations) in Nikon Elements software for dendritic spine analysis.

**Imaris Volumetric Reconstruction**

*CD68 and CD11b:* A “surface” module was applied to each image in which the threshold absolute intensity was selected to apply the surface rendering and quantify the volume of immunostaining. The threshold value was selected by fine-tuning the automatic detection value to accurately detect positive staining in each image. A value was generated per image based on the total immunoreactivity volume (μm^3^) rendered in the image.

*Iba1:* A “filaments” module was applied to each image and settings were consistent across images: largest diameter (9 μm), thinnest point (0.6 μm), diameter of sphere region (10 μm), seed point threshold (500), and dendrite diameter threshold (2.5). The starting seed points were manually selected for individual microglia (the whole process arbor was intact and within image borders) and placed at the center of the soma. Non-optimal microglia where also seeded to reduce the number of false/continuous tracings. Once rendered, microglial values were sorted by volume (sum) and 8-12 whole microglia per image were selected and Sholl, volume (sum), and branch points were measured.

*Thy1:* Due to inconsistencies in Imaris 3D reconstruction tracing and spine detection between images when visually inspected, Thy1-YFP labeled dendritic spines were manually counted using the Golgi counting method (5).

**Microglial Sholl Analysis**

In addition to changes in the expression of activation markers, microglia display dramatic morphological changes that include both extension and retraction of processes as they rapidly respond to fluctuations in the microenvironment and converge toward sites of injury. Over time, in chronic injuries, microglial morphology shifts from a surveying, ramified morphology to an activated, more amoeboid profile (6,7). Microglial morphology can be visualized through Iba1 staining and quantified via Sholl analysis to display the process arbor as a topographical map (8). To further capture the complexity of the microglial arbor in 3D space, we used a volumetric Sholl analysis to quantify the number of intersections as a function of spherical distance (2 μm steps) from the cell soma. Our data demonstrate that WT male or KO mice did not display an irradiation-mediated effect while WT females displayed a significant loss in medial processes in response to radiation (Supplemental Fig. 4a – WT female: F_(23, 192)_ = 2.012, p = 0.0058, *post hoc* 14-24 μm, p < 0.05; Supplemental Fig. 4c – WT female: p = 0.0496). Interestingly, while the interaction between irradiation and process intersection was not significant (Supplemental Fig. 4b – KO male: F_(23, 192)_ = 1.138, p = 0.3075, *post hoc* 26-40 μm, p < 0.05) there was a trend towards increased distal processes in KO male mice. Further, while there were no overt sex differences in WT mice, sham KO mice demonstrated significant differences in baseline distal processes intersections (Supplemental Fig. 4b – sham KO: F_(23, 168)_ = 7.451, p < 0.0001, *post hoc* 12-42 μm, p < 0.05 ) and area under the curve (Supplemental Fig. 4c – sham KO: p = 0.0365). Overall, these results demonstrate subtle changes to microglia morphology and suggest that a single 10 Gy dose is not sufficient to cause lasting changes in morphology 45 d post-irradiation.

Microglial morphology was also comparable in vehicle and LA1 treated sham mice; though, following irradiation, vehicle treated microglia had reduced distal processes while LA1 treatment resulted in increased processes (Supplemental Fig. 4e – non-significant interaction, two-way ANOVA: F_(23, 192)_ = 0.6935, p = 0.9679, *post hoc* 20-32 μm, p < 0.05). Although this difference was subtle, it provides evidence for a differential irradiation response where LA1 treated microglia display an increased arbor complexity that may be indicative of a dampened activation profile.

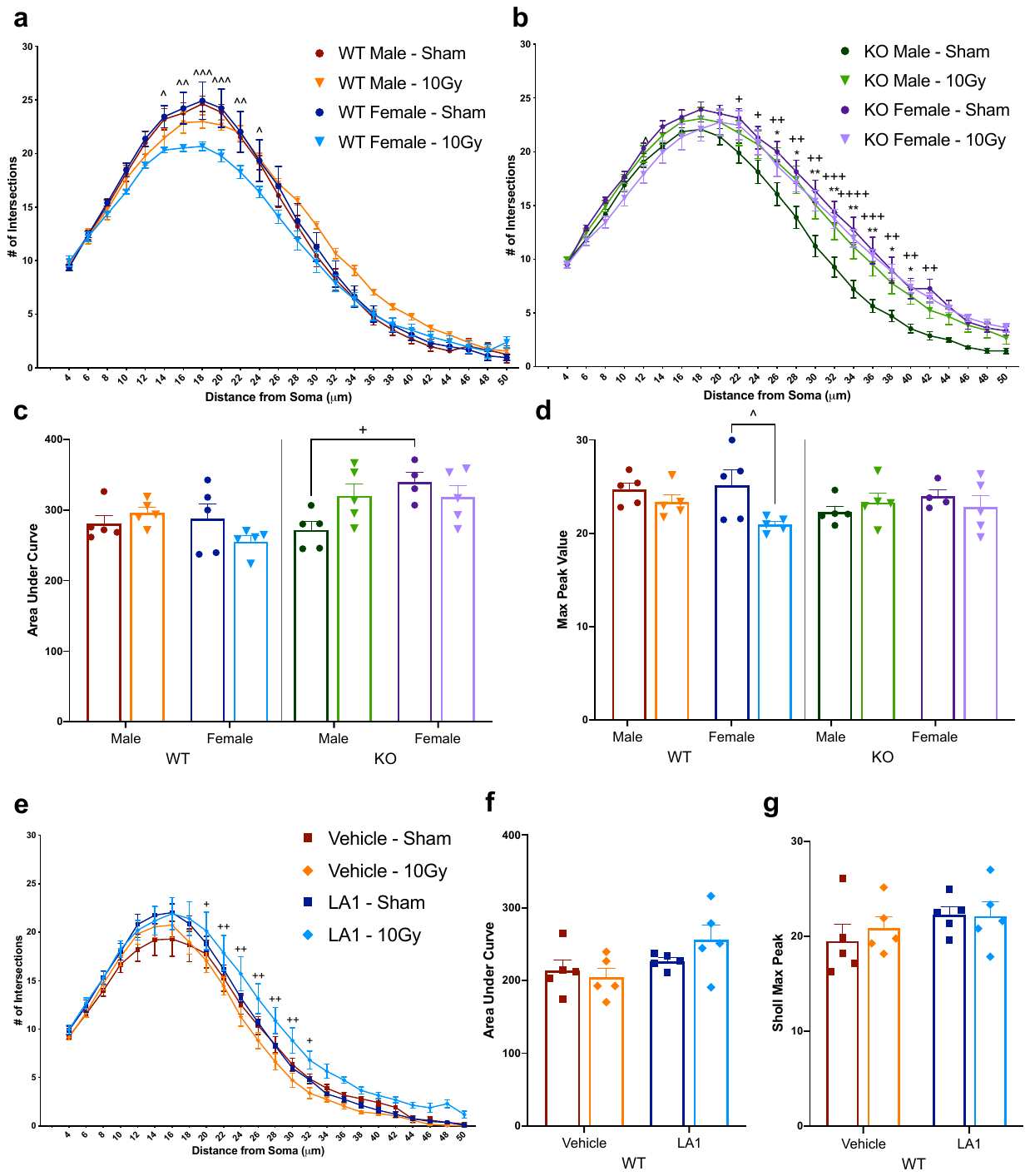

**Supplemental Figure 4. Morphological volumetric Sholl analysis of Iba1 stained microglial arbor** showing **a**) WT mice and **b**) KO mice. To further demonstrate differences **c**) area under the curve and **d**) max peak values were plotted demonstrating that irradiation had a significant effect on female WT medial processes. Further, distal processes were affected in KO animals and differed between sexes. n = 5 per group; **a, b**) two-way ANOVA with multiple comparisons per step; **c, d**) three-way ANOVA followed by multiple comparisons test. * p< 0.05, ** p< 0.01, *** p<0.001, **** p<0.0001. Sham vs irradiated significance: * male, ^ female. + male vs female sham significance.
